# Heteroallelic combination of anillin mutants reveals cytoskeletal uncoupling during cytokinesis

**DOI:** 10.1101/2024.04.21.590302

**Authors:** Guillaume Chambaud, Sabrya C. Carim, Gilles R. X. Hickson

## Abstract

Cytokinetic contractile rings close, disassemble and transition to midbody rings through poorly understood mechanisms. We previously showed that this transition in *Drosophila* S2 cells requires both Citron kinase (Sticky) and septins, acting on the N- and C-terminus, respectively, of the scaffold protein, Anillin. To better understand the coordination of these proteins, we performed deplete-and-rescue experiments using Anillin mutants defective in Sticky interaction or septin recruitment. Expressing each mutant individually as two copies tagged with GFP and mCherry revealed comparable behaviors of the tags and characteristic cytokinesis failure phenotypes. However, trans-heterozygous combination of both Anillin mutants rescued the cytokinesis defects, with both mutants clearly exhibiting different localization patterns. Thus, the essential N- and C-terminus-mediated roles of Anillin can be satisfied by different molecules of Anillin that cannot bind to Sticky and to septins simultaneously. We conclude that Anillin organizes distinct cytoskeletal elements that are uncoupled from one another and that their physical separation drives the transition from contractile ring to midbody ring.

## Introduction

The metazoan cytokinetic contractile ring (CR) forms in response to RhoA activation at the equatorial plasma membrane and through the subsequent recruitment of many proteins including formins, the RhoA-dependent kinases, Rho-kinase and Citron kinase, actin filaments, myosin II motor proteins, septins and the scaffold protein Anillin (D’Avino et al., 2015; Glotzer, 2017; Green et al., 2012; Pollard and O’Shaughnessy, 2019). Through poorly understood mechanisms dependent upon actomyosin contractility (Pollard, 2017), the membrane anchored CR then constricts, reeling material in from its flanks, via cortical flow, while somehow simultaneously shrinking in circumference and disassembling (Schroeder, 1972). Upon reaching a diameter of approximately 1 µm, and upon encircling the microtubules of the midbody, the CR transitions to a more stable midbody ring (MR) structure. The MR firmly anchors the plasma membrane until the later events of abscission complete cytokinesis, except in cases of developmentally programmed incomplete cytokinesis wherein the MR persists as a stable ring canal (Gerhold et al., 2022; Haglund et al., 2011; Price et al., 2023). The transition from CR to MR is a crucial stage of the cell cycle necessary for tissue homeostasis and to prevent the formation of binucleate tetraploid cells, which are prone to form cancers (Fujiwara et al., 2005; Ganem et al., 2007).

How different cytoskeletal components are organized and reorganized during closure of the CR and formation of the MR is unclear, but Anillin is heavily implicated (Carim et al., 2020; D’Avino, 2009; Hickson and O’Farrell, 2008a; Piekny and Maddox, 2010). Anillin is a scaffold for actomyosin (Field and Alberts, 1995; Jananji et al., 2017; Kučera et al., 2021; Straight et al., 2005; Tian et al., 2015), a mediator of septin recruitment (Carim and Hickson, 2023; Hickson and O’Farrell, 2008b; Liu et al., 2012; Maddox et al., 2005), and a protein required for MR biogenesis and maturation (El Amine et al., 2013; Kechad et al., 2012; Panagiotou et al., 2022; Renshaw et al., 2014). Anillin also binds Citron kinase (Sticky in *Drosophila*), a RhoA target protein which, although recruited early to CRs, is specifically required later for formation of a stable MR (Bassi et al., 2011; D’Avino et al., 2004; Echard et al., 2004; El-Amine et al., 2019; Gai et al., 2011; Price et al., 2023). Numerous other interactors of Anillin have been reported, including RacGAP50C (D’Avino et al., 2008), ECT2 (Frenette et al., 2012), Cindr/CD2AP (Haglund et al., 2010), Syndapin (Takeda et al., 2013), importins (Beaudet et al., 2020). Important outstanding questions relate to how Anillin’s interactions are coordinated. Does Anillin bind its interactors simultaneously and link them together? Or does Anillin form mutually exclusive complexes with different partners, perhaps sequentially in a dynamic cycle, or at different stages of cytokinetic progression? As a central and essential component of the cytokinetic machinery, understanding Anillin function promises to unlock the longstanding mystery of how that machinery operates.

We previously reported that opposing actions of Sticky, acting via the N-terminus of Anillin, and septins, acting via the C-terminus of Anillin, are required for the transition from CR to MR of *Drosophila* S2 cells (El Amine et al., 2013). This conclusion was based primarily on the following observations: 1) membrane-associated Anillin and septins are ordinarily shed from the nascent MR as it thins to become a stable, mature MR; 2) depletion of the septin Peanut blocked the shedding of Anillin, but not stable MR formation; 3) Sticky depletion reciprocally blocked stable MR formation, but not shedding; 4) Sticky-dependent retention of Anillin required the Anillin N-terminus, while Peanut-dependent shedding required the Anillin C-terminus (El Amine et al., 2013). These observations led us to propose a tug-of-war model in which Anillin connects both Sticky and septins at the same time, with the former promoting Anillin retention in ring structures and the latter promoting Anillin’s removal (El Amine et al., 2013). More recently, we have shown that Sticky is a *bona fide* Rho1-dependent protein that interacts with the Anillin N-terminal domain, as does mammalian Citron kinase (Gai et al., 2011), and that both the Anillin N-terminal domain and Sticky’s interaction with Rho1-GTP are required for stable MR formation (El-Amine et al., 2019). We have also shown that the ability of the Anillin C-terminal PH domain to recruit septins to the CR (Field et al., 2005; Liu et al., 2012; Oegema et al., 2000) specifically requires Anillin’s interaction with Rho1-GTP (Carim and Hickson, 2023), shown by others to occur via the C-terminal AH domain (Piekny and Glotzer, 2008; Sun et al., 2015). In the present study, we sought to better understand how Sticky, Anillin and septins collaborate during the crucial transition from CR closure to MR formation. To this end, we have exploited separation-of-function alleles of Anillin that perturb either interaction with Sticky, or the recruitment of septins, each with severe consequences to the success of cytokinesis. Remarkably, we find that, when co-expressed in a trans-heterozygous combination, these alleles complement one another to rescue cytokinesis. Rather than supporting a tug-of-war model in which Anillin links Sticky and septins together, the data provide compelling evidence that furrow-associated Anillin molecules act either with Sticky or with septins in a mutually exclusive manner. This provides key mechanistic insight into how CR closure and disassembly are coordinated with the formation of a stable MR, essential processes in the division of animal cells.

## Results

### A novel separation-of-function allele, Anillin-Δ1-5, fails to interact with Sticky and fails to support a faithful CR-to-MR transition

We previously found that *Drosophila* Anillin (the product of the *scraps* gene) and Sticky interact via their NTD and mini-CC2a regions, respectively (see Fig. 1A for domain organization of Anillin and Sticky), and that deletion of the Anillin NTD prevented the interaction and disrupted MR formation (El-Amine et al., 2019). However, the Anillin NTD comprises 150 amino acids, which could potentially also interact with other components such as Cindr (Haglund et al., 2010), or Diaphanous, the homolog of mammalian mDia2 which interacts with the ANLN NTD (Watanabe et al., 2010). We therefore wished to generate a more specific separation-of-function allele of Anillin that cannot interact with Sticky. We noted that, while the sequence of the Anillin NTD has not been particularly well conserved through evolution, the first 5 amino acids are identical from *Drosophila* to humans (Fig. 1B). Alphafold-2-predicted structures of metazoan anillins all contain 18-24 residue alpha-helices at their extreme N-termini, which otherwise appear to be largely unstructured (see Fig. S1 for predicted structure of *Drosophila* Anillin). We hypothesized that deletion of the highly conserved first 5 amino acids would abrogate the interaction with Sticky. Indeed, unlike the GST-Anillin-NTD positive control, a recombinant GST-Anillin-NTDΔ1-5 failed to pull down Sticky-miniCC2a-GFP from S2 cell lysates (Fig. 1C and Fig. S1 for full blots). Similarly, unlike wild-type Anillin, but similar to Anillin-ΔNTD, Anillin-Δ1-5 was unable to recruit Sticky mini-CC2a to the CR/nascent MR (Fig. 1D). We conclude that the conserved, first 5 amino acids at the extreme N-terminus of *Drosophila* Anillin are required for the interaction between Anillin and Sticky.

**Fig. 1.**
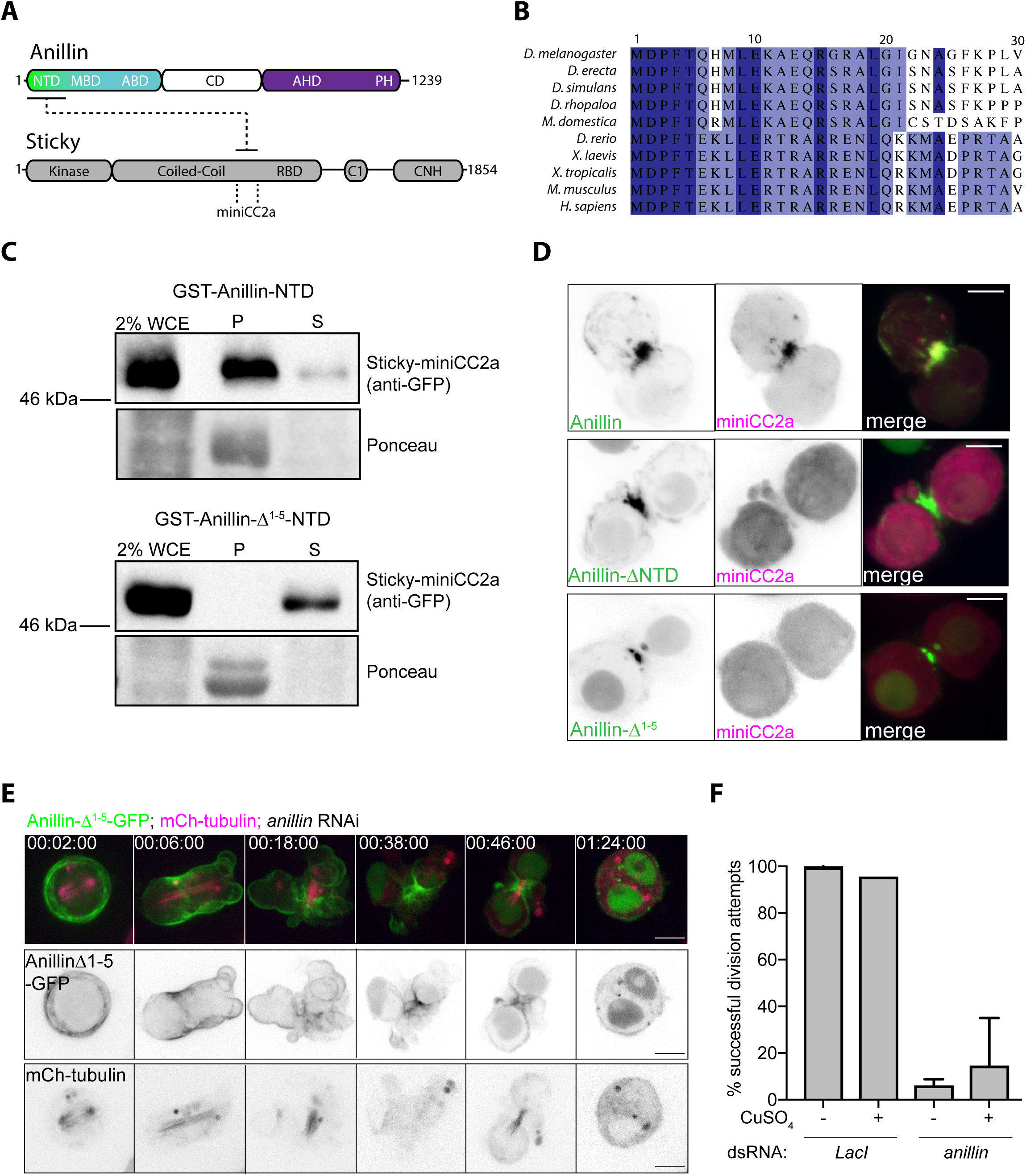
A novel separation-of-function allele, Anillin-Δ1-5, fails to interact with Sticky and fails to support a faithful CR-to-MR transition. **A** Domain organization of *Drosophila* Anillin and Sticky, including the mini-CC2a region of Sticky previously shown to interact with the Anillin N-terminal domain (NTD). MBD, myosin binding domain; ABD, actin binding domain; CD, central domain; RBD, Rho1 binding domain; AHD, Anillin homology domain; PH, Pleckstrin homology domain; CNH, Citron-Nik1-homology domain. **B** Sequence alignment of the extreme N-terminus of Anillin from various species. **C** Pulldowns of Sticky-miniCC2a-GFP expressed in S2 cells by GST-Anillin-NTD (upper blots) or GST-Anillin-Δ1-5-NTD (lower blots), stained with Ponceau-S or anti-GFP antibody. P, pellet; S, supernatant; WCE, whole cell extract. Note the loss from the pellet fraction of Sticky-miniCC2a-GFP in the GST-Anillin-Δ1-5-NTD pulldown. Full blots are shown in Fig. S1. **D** Still images from live-cell recordings of S2 cells depleted of endogenous Anillin and co-expressing miniCC2a-mCherry and Anillin-GFP (upper panel), Anillin-ΔNTD-GFP (middle panel), or Anillin-Δ1-5-GFP (lower panel). Note that only wild-type Anillin-GFP can recruit miniCC2a-mCherry to the division plane. **E** Images from a representative time-lapse sequence of an S2 cell constitutively expressing mCherry-tubulin and inducibly expressing Anillin-Δ1-5-GFP, following 5 days of *anillin* (*scraps*) RNAi (see Video 1). Times are h: min: sec; scale bars, 5 µm. **F** Quantification of successful division attempts in cells transiently transfected and induced to express (or not, with CuSO_4_) Anillin-Δ1-5-GFP, following 5 days RNAi of LacI (negative control) or endogenous Anillin. Mean ± SD from n=2 independent experiments shown.

We examined the functionality of Anillin-Δ1-5 in live-cell, deplete-and-rescue experiments (Fig. 1E-F). Induced expression of RNAi-resistant Anillin-Δ1-5-GFP in transiently transfected S2 cells failed to rescue cytokinesis following RNAi-mediated depletion of endogenous Anillin. Furrowing proceeded relatively normally, albeit with pronounced blebbing of the plasma membrane (see Video 1), but 86% of division attempts failed at the MR stage (Fig. 1E-F). These results both support the prior conclusion that the Anillin-Sticky interaction is essential for the CR-to-MR transition (El-Amine et al., 2019) and validate Anillin-Δ1-5 as a separation-of-function allele.

We also monitored Sticky-GFP localization in cells expressing Anillin-Δ1-5-mCherry and depleted of both endogenous proteins. When co-expressed with wild-type Anillin- mCherry, Sticky-GFP was robustly recruited to the CR and MR (Fig. S1). However, when co-expressed with Anillin-Δ1-5-mCherry, Sticky-GFP localization was significantly reduced, both at the furrow cortex and at the nascent MR (Fig. S1). These data further support the conclusion that Anillin-Δ1-5 lacks the ability to interact with Sticky and suggest that the interaction with Anillin enhances the recruitment and/or retention of Sticky.

### GFP- and mCherry-tagged versions of Anillin behave similarly when co-expressed

We recently reported the phenotype induced by another Anillin mutant, Anillin- A874D/E829K (hereafter referred to as Anillin-RBD* as in (Carim and Hickson, 2023)), which harbors two amino acid substitutions in the Rho1 binding domain (RBD) that prevent binding to Rho1-GTP (Sun et al., 2015). Anillin-RBD* failed to recruit septins to the CR, failed to promote timely furrowing, and failed to be shed during the transition to the MR (Carim and Hickson, 2023). However, despite highly penetrant cytokinesis failures, prominent,

Sticky-positive, MR-like structures still formed in the presence of Anillin-RBD*, suggesting that Anillin-RBD* retained the ability to interact with Sticky (Carim and Hickson, 2023).

To better understand the functional relationships between Anillin, Sticky and Rho1/septins, we wished to examine the consequences of Anillin-Δ1-5 and Anillin-RBD* mutations together, both as a double mutant (in *cis*), but also as a *trans*-heterozygous combination, with each allele fused to either GFP or mCherry and co-expressed in the same cells. To validate this approach, we first performed control experiments in which we co-expressed wild-type Anillin-GFP and Anillin-mCherry in transiently transfected S2 cells depleted of endogenous Anillin. This revealed extensive co-localization of GFP and mCherry throughout cytokinetic progression, as expected (Fig. 2A and Video 2). From time-lapse sequences, we quantified the cortical levels of GFP and mCherry, by obtaining line profile measurements along the cortex from one pole to the other for each dividing cell (Fig. 2B). The sum intensities along each line increased over time as the cells underwent cytokinesis, and similarly for both GFP and mCherry channels, indicating a global recruitment of both Anillin species at the cortex (Fig. 2C). For each timepoint, the measurement line was also sub-divided into four segments of equal length: the outer two segments designated as “polar” and the central two as “equatorial”. For each channel, the sum of pixel intensities of the “equatorial” and “polar” segments of the line were then calculated and expressed as equatorial: polar ratios (Fig. 2D). As expected, this revealed that, as the cells underwent cytokinesis, cortical Anillin became progressively enriched at the equator, relative to the polar cortex. Importantly, the degree of enrichment measured was comparable for both Anillin-GFP and Anillin-mCherry. These Anillin-GFP and Anillin-mCherry co-expressing cells closed their CRs within 12 minutes of anaphase onset, showed clear evidence of normal shedding from the nascent MRs, and faithfully underwent the transition to stable and mature MRs that did not regress to form binucleate cells (Fig. 2A).

**Fig. 2.**
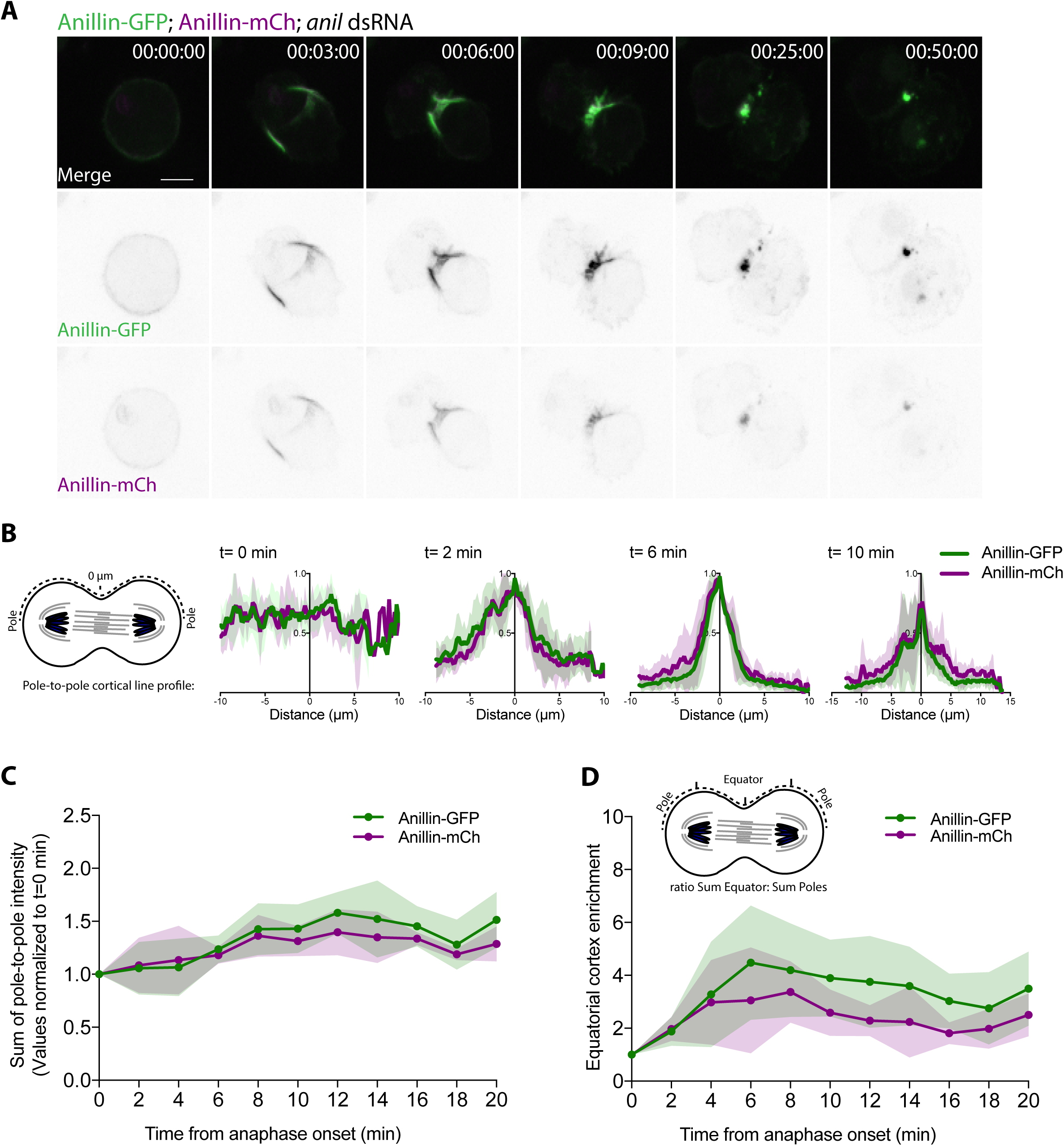
GFP- and mCherry-tagged versions of Anillin behave similarly when co-expressed. **A** Images from a representative time-lapse sequence of an S2 cell co-expressing Anillin-GFP (green in upper merged panel, inverted LUT in middle panel) and Anillin-mCherry (magenta in merged, inverted LUT in lower panel), following RNAi-mediated depletion of endogenous Anillin (see Video 2). Times are h: min: sec from anaphase onset; scale bars, 5 µm. **B** Intensity profiles of Anillin-GFP and Anillin-mCherry obtained from lines drawn, from pole to pole, along the cortex of individual cells at successive times from anaphase onset (T=0) as they progress through furrowing. For each line and each channel, background-corrected intensity values were normalized to the highest value along the line to show the relative distributions of signal for each channel along the cortex (mean ± SD, N= 30). **C** Quantification of cortical enrichment for each channel, calculated as the sum of all intensity values per channel per line, plotted through time and normalized to the values at T=0. Mean values ± SD are shown for each channel. **D** Quantification of equatorial enrichment for each channel, calculated as the fold increase of sum intensity at the equatorial cortex (defined as the two central 25% lengths of each line) relative to the sum intensity at the polar cortex (defined as the two outer 25% lengths of each line). Mean values ± SD are shown for each channel.

### Co-expression of GFP- and mCherry-tagged versions of each single Anillin mutant faithfully reproduces phenotypes

We next examined cells transiently co-expressing Anillin-Δ1-5-GFP and Anillin-Δ1-5- mCherry in cells depleted of endogenous Anillin for 3 days (Fig. 3A-D and Video 3). Once again, extensive colocalization of GFP- and mCherry was observed and the phenotypes resembled those observed upon expression of Anillin-Δ1-5-GFP alone (Fig. 3A). From 3 independent experiments, 58 of 97 observed division attempts (i.e. 60%) ended in failed cytokinesis (binucleation), with a mean time of failure 257 min after anaphase onset (Fig. 3B). A slight, non-significant delay in the speed of CR closure was observed, compared to the wild-type controls co-expressing Anillin-GFP and Anillin-mCherry (Fig. 3C). The equatorial cortex enrichment profiles of co-expressed Anillin-Δ1-5-GFP and Anillin-Δ1-5-mCherry were also measured and found to be comparable to one another (Fig. 3D).

**Fig. 3.**
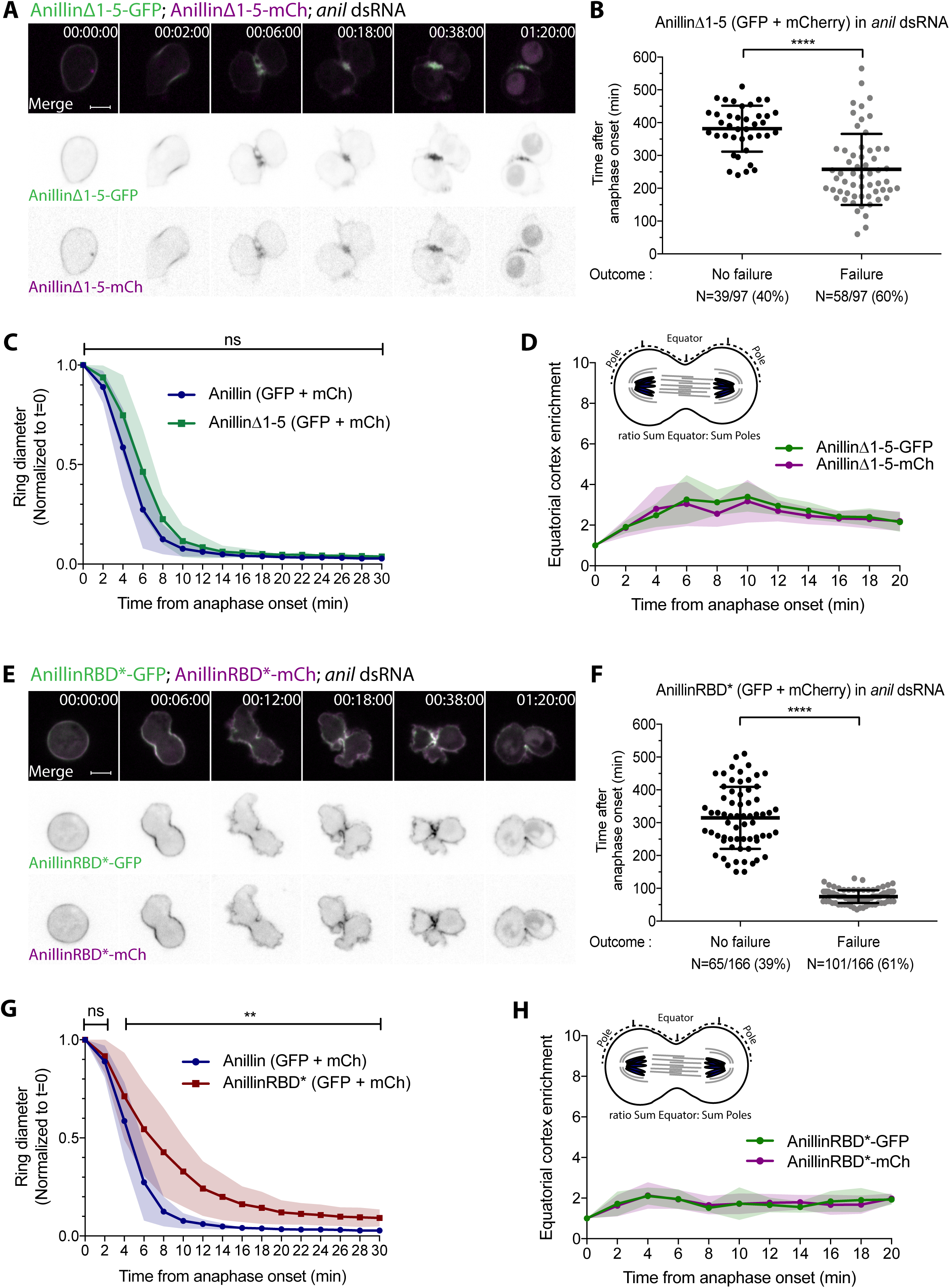
GFP- and mCherry-tagged versions of each single Anillin mutant faithfully reproduce phenotypes when co-expressed. **A** Stills from a time-lapse sequence of a representative S2 cell co-expressing Anillin-Δ1-5-GFP (green in merged, inverted LUT in middle panel) and Anillin-Δ1-5-mCherry (magenta in merged, inverted LUT in lower panel), following RNAi-mediated depletion of endogenous Anillin (see Video 3). **B** Outcomes of 97 division attempts imaged and the timing of stated outcome relative to anaphase onset. Note that “failure” corresponds to observed furrow regression and binucleation, while “no failure” can signify either successful abscission or the end of the recording, and thus may include cells that were destined to fail. **C** Contractile ring diameter measurements through time, normalized to the cell diameter at anaphase onset (T=0). Mean values ± SD are shown for cells expressing Anillin-Δ1-5 (-GFP/-mCherry, N=28) as well as for wild-type Anillin (-GFP/-mCherry, N=30), in cells depleted of endogenous Anillin for 3 days. **D** Quantification of equatorial cortex enrichment for each channel, calculated as the fold increase of sum intensity along the equatorial cortex (defined as the two central 25% lengths of each line) relative to the sum intensity along the polar cortex (defined as the two outer 25% lengths of each line), N=25 cells. **E** Stills from a representative time-lapse sequence of an S2 cell co-expressing Anillin-RBD*-GFP (green in merged, inverted LUT in middle panel) and Anillin-RBD*-mCherry (magenta in merged, inverted LUT in lower panel), following 3-day RNAi-mediated depletion of endogenous Anillin (see Video 4). **F** Outcomes of 166 division attempts imaged and the timing of stated outcome relative to anaphase onset. Note that “failure” corresponds to observed furrow regression and binucleation, while “no failure” can signify either successful abscission or the end of the recording, and thus may include cells that were destined to fail. **G** Contractile ring diameter measurements through time, normalized to the cell diameter at anaphase onset (T=0). Mean values ± SD are shown for cells expressing Anillin-RBD* (-GFP/-mCherry, N=32) as well as for wild-type Anillin (-GFP/-mCherry), in cells depleted of endogenous Anillin for 3 days. **H** Quantification of equatorial cortex enrichment for each channel, calculated as the fold increase of the sum intensity along the equatorial cortex (defined as the two central 25% lengths of each line) relative to the sum intensity along the polar cortex (defined as the two outer 25% lengths of each line), N=30 cells. Times are h: min: s from anaphase onset; ns, not significant; *, p>0.05; **, p<0.01; ****, p<0.0001; scale bars, 5 µm.

We next examined cells transiently co-expressing Anillin-RBD*-GFP and Anillin- RBD*-mCherry in cells depleted of endogenous Anillin (Fig. 3E-H). Once again, GFP and mCherry colocalized extensively (Fig. 3E and Video 4) and the phenotypes were comparable to those reported for Anillin-RBD*-GFP expression alone (Carim and Hickson, 2023). These included failed cytokinesis (binucleation of 101 of 166 division attempts, Fig. 3F), significantly slowed CR closure (Fig. 3G), the absence of detectable shedding (Fig. 3E), and the formation of internal MR-like structures (Fig. 3). The equatorial cortex enrichment profiles of co-expressed Anillin-RBD*-GFP and Anillin-RBD*-mCherry were also very similar to one another (Fig. 3H). Together, these experiments validate the approach of comparing GFP- and mCherry-tagged versions of Anillin, which behave similarly when co-expressed in the same cells, and which produce similar phenotypes to those observed when only a single construct is expressed.

### An Anillin-Δ1-5-RBD* double mutant is severely defective, but still retains some function

We hypothesized that a double mutant (in *cis*) of Anillin that is defective in both Sticky and Rho1 binding would be severely hampered in its ability to localize and function. Co-expression of GFP- and mCherry-tagged versions of such a double mutant is shown in Fig. 4A and Video 5. Despite sill harboring myosin- and actin-binding domains, Anillin-Δ1-5-RBD* was only barely detectable at the furrow cortex at the CR stage. As the cells attempted cytokinesis, the cortex exhibited many blebs in both equatorial and polar regions. Blebs are indicative of force imbalances and/or loss of attachment between the cortex and the plasma membrane. These blebs were more frequent and more pronounced than any observed with the Anillin-Δ1-5 or Anillin-RBD* single mutants alone, suggesting that interactions between Anillin and Sticky, and Anillin and Rho1/septins, each ordinarily act to suppress blebs during furrowing. The presence of such pronounced blebs in Anillin-Δ1-5-RBD* cells impeded our ability to reliably trace the cortex from pole to pole and quantify the total cortical equatorial versus polar pools of Anillin. However, measurements of the cortex to cytoplasm enrichment from line profiles across the equator were similar for both GFP- and mCherry-tagged Anillin-Δ1-5-RBD* double mutants, and markedly diminished when compared to co-expressed wild-type Anillin (Fig. S2). Aberrant Anillin-Δ1-5-RBD*-dependent CRs closed slowly and incompletely (Fig. 4B) and, following 3 days of RNAi, resulted in 63% of division attempts failing cytokinesis (77 out of 122 observed, Fig. S2). In cells that were destined to fail, Anillin-Δ1-5-RBD* was still recruited to nascent MR-like structures that appeared later during the abortive furrowing (Fig. 4A). These MR-like structures did not exhibit shedding and failed to anchor the plasma membrane, which ultimately regressed. These results demonstrate that the Anillin-Δ1-5-RBD* double mutant is at least as perturbing as each of the single mutants, while still retaining some function.

**Fig. 4.**
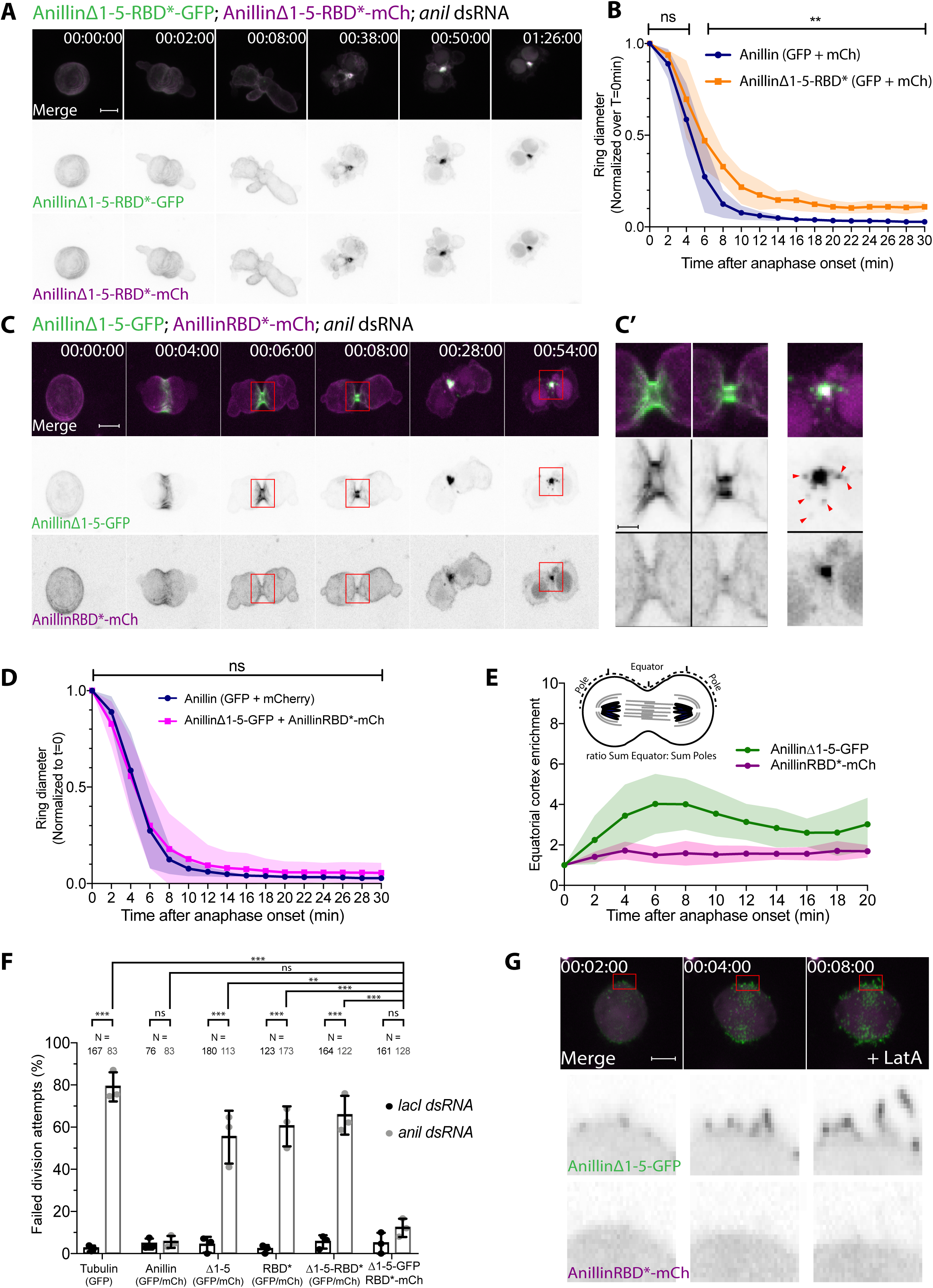
An Anillin-Δ1-5-RBD* double mutant is severely defective, while the trans-heterozygous allelic combination rescues cytokinesis. **A** Stills from a representative time-lapse sequence of an S2 cell co-expressing double mutant Anillin-Δ1-5-RBD* fused to GFP (green in merged, inverted LUT in middle panel) or mCherry (magenta in merged, inverted LUT in lower panel), following 3-day RNAi-mediated depletion of endogenous Anillin (see Video 5, and Fig. S2 for quantification of outcomes). **B** Contractile ring diameter measurements through time, normalized to the cell diameter at anaphase onset (T=0). Mean values ± SD (N=27) are shown for cells co-expressing Anillin-Δ1-5-RBD* (-GFP/-mCherry) as well as for wild-type Anillin (-GFP/-mCherry), in cells depleted of endogenous Anillin. **C** Stills from a time-lapse sequence of a representative S2 cell co- expressing the trans-heterozygous combination of Anillin-Δ1-5-GFP (green in merged, inverted LUT in middle panel) and Anillin-RBD*-mCherry (magenta in merged, inverted LUT in lower panel), following RNAi-mediated depletion of endogenous Anillin (see Video 6). **C’** Zoomed views of the boxed regions in C. Arrowheads point to shed particles apparent in the GFP channel only. **D** Contractile ring diameter measurements through time, normalized to the cell diameter at anaphase onset (T=0). Mean values ± SD (N=30) are shown for cells expressing Anillin-Δ1-5-GFP and Anillin-RBD*-mCherry, as well as for wild-type Anillin (- GFP/-mCherry), in cells depleted of endogenous Anillin. **E** Quantification of equatorial cortex enrichment of co-expressed Anillin-Δ1-5-GFP and Anillin-RBD*-mCherry, calculated as the fold increase of sum intensity at the equatorial cortex (defined as the two central 25% lengths of each line) relative to the sum intensity at the polar cortex (defined as the two outer 25% lengths of each line), N=30 cells. **F** Quantification of failed division attempts (i.e. furrow regression and binucleation) observed from live recordings of cells expressing the indicated constructs, following incubation with lacI (control) or anillin dsRNAs. **G** Stills from a time- lapse sequence of a representative S2 cell, depleted of endogenous Anillin, and co-expressing Anillin-Δ1-5-GFP (green in merged, inverted LUT in middle panel) and Anillin-RBD*-mCherry (magenta in merged, inverted LUT in lower panel), attempting cytokinesis in the presence of 1 µg/ml LatA (see Video 7). Times are h: min: s from anaphase onset; ns, not significant; *, p>0.05; **, p<0.01; ****, p<0.0001; scale bars, 5 µm.

### Co-expression of Anillin-RBD* and Anillin-Δ1-5 in a trans-heterozygous combination rescues cytokinesis

Next, we examined the trans-heteroallelic combination of Anillin-Δ1-5-GFP and Anillin-RBD*-mCherry in cells depleted of endogenous Anillin (Fig. 4C and Video 6). Remarkably, this restored the normal speed of CR closure (Fig. 4D) and largely rescued the failed cytokinesis induced by each mutant alone (Fig 4F). Quantitative analysis of cortical line profiles revealed a more pronounced total cortical recruitment of Anillin-Δ1-5 over time than co-expressed Anillin-RBD* (Fig. 4E). Anillin-Δ1-5 was also more prominently recruited earlier, and became more equatorially enriched, with a mean equator: pole ratio of approximately 4, compared to Anillin-RBD*, which remained more broadly distributed along the cortex, with a mean equator: pole ratio of approximately 2 (Fig. 4E). The localization patterns of the two co-expressed proteins were also clearly different (Fig. 4C). Following its initial robust recruitment, Anillin-Δ1-5 showed clear evidence of shedding from the nascent MR (see arrowheads in Fig. 4C’). Conversely, Anillin-RBD* was only weakly recruited initially, then accumulated at the nascent MR, where it was retained without showing any evidence of shedding (Fig. 4C and 4C’).

Dividing cells were also pre-treated with 1 µg/ml Latrunculin A (LatA), which prevents F-actin polymerization but does not prevent the Rho1-dependent recruitment of Anillin and septins to the plasma membrane (Hickson and O’Farrell, 2008b). In LatA, Anillin-Δ1-5 localized to the plasma membrane in similar fashion to wild-type Anillin, while co-expressed Anillin-RBD* remained entirely cytoplasmic (Fig. 4G and Video 7). Thus, in both the presence and absence of F-actin, co-expressed Anillin-Δ1-5 and Anillin-RBD* exhibited clearly distinct localization patterns and behaved independently of one another. Yet, in the presence of F-actin, these two alleles were able to functionally complement one another. To confirm that the observed differences were not due to the GFP/mCherry tags, or our abilities to detect them, we swapped them. Co-expression of Anillin-Δ1-5-mCherry and Anillin-RBD*- GFP produced comparable results (Fig. S3).

Altogether, these results clearly reveal that the cytokinetic apparatus can successfully utilize and separate distinct pools of Rho1-binding-defective and Sticky-binding-defective Anillin. We conclude that individual Anillin molecules do not need to bind to Sticky and to Rho1/septins at the same time. Accordingly, we propose that the normal Sticky-dependent retention versus septin-dependent removal of Anillin previously reported (El Amine et al., 2013), reflects the physical separation of distinct Anillin-scaffolded cytoskeletal elements that are independently anchored to the plasma membrane.

### Mutant alleles exhibit differential abilities to compete with wild-type Anillin for equatorial localization

Finally, we exploited this co-expression system to compare the abilities of mCherry-tagged mutant alleles to compete with wild-type Anillin-GFP for localization. Specifically, we compared the equatorial cortex-to-cytoplasm enrichment of the two channels, each measured from the same line per cell as it progressed through anaphase and cytokinesis. Anillin-Δ1-5-mCherry exhibited an equatorial cortex enrichment profile through time that was similar to co-expressed wild-type Anillin-GFP, confirming that the interaction with Sticky is not a major contributor to the furrow localization pattern of Anillin (Fig. 5A-B and Video 8). Conversely, Anillin-RBD*-mCherry showed a markedly reduced ability, compared to co-expressed wild-type Anillin-GFP, to become enriched at the equatorial cortex during anaphase (Fig. 5C-D and Video 9). This confirms that Rho1-GTP binding (and subsequent septin binding as shown in (Carim and Hickson, 2023)) does provide a major contribution to the total levels of Anillin at the equatorial cortex during anaphase.

**Fig. 5.**
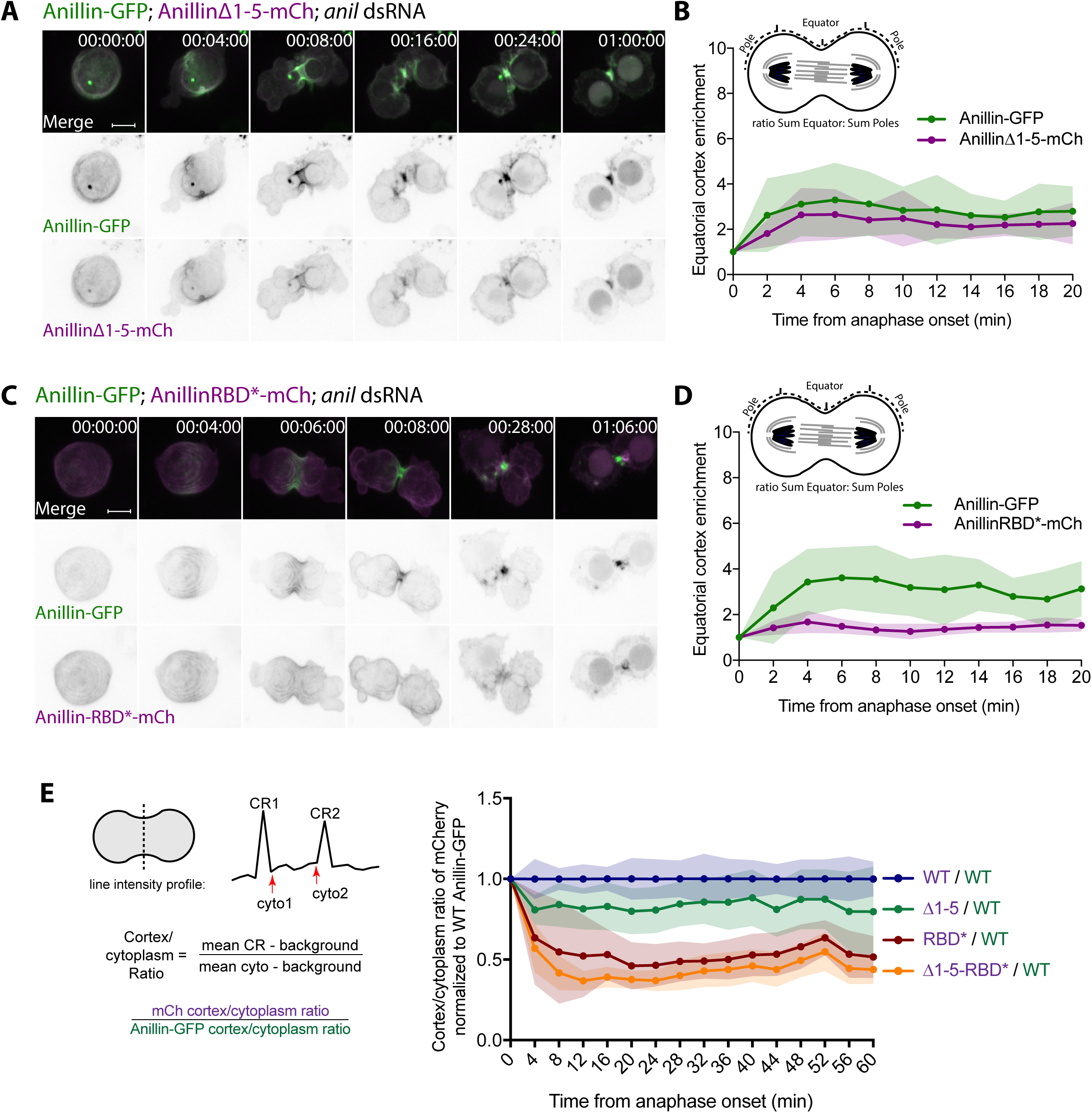
Differential abilities of the mutant alleles to compete with wild-type Anillin for equatorial localization. **A** Stills from a time-lapse sequence of a representative S2 cell co-expressing wild-type Anillin-GFP (green in merged, inverted LUT in middle panel) and Anillin-Δ1-5-mCherry (magenta in merged, inverted LUT in lower panel), following RNAi-mediated depletion of endogenous Anillin (see Video 8). **B** Quantification of equatorial cortex enrichment for each channel, calculated as the fold increase of sum intensity at the equatorial cortex (defined as the two central 25% lengths of each pole-to-pole line) relative to the sum intensity at the polar cortex (defined as the two outer 25% lengths of each line), N=23 cells. **C** Stills from a time-lapse sequence of a representative S2 cell co-expressing wild-type Anillin-GFP (green in merged, inverted LUT in middle panel) and Anillin-RBD*-mCherry (magenta in merged, inverted LUT in lower panel), following RNAi-mediated depletion of endogenous Anillin (see Video 9). **D** Quantification of equatorial cortex enrichment for each channel, calculated as the fold increase of sum intensity at the equatorial cortex (defined as the two central 25% lengths of each pole-to-pole line) relative to the sum intensity at the polar cortex (defined as the two outer 25% lengths of each line). **E** The cortex: cytoplasm ratios of the indicated mCherry- tagged constructs each normalized to the cortex: cytoplasm ratio of wild-type Anillin-GFP co- expressed in the same cell. This reveals the relative ability of each mutant to compete with wild-type Anillin for localization to the equatorial cortex.

We also calculated the cortex: cytoplasm ratios of each mCherry-tagged mutant and normalized it to the equivalent ratio of co-expressed Anillin-GFP (Fig. 5E). This produced a ratio of 1.0 for control cells co-expressing Anillin-mCherry and Anillin-GFP, while ratios fell below 1.0 for the other combinations, in the following order: Anillin-Δ1-5, Anillin-RBD*, Anillin-Δ1-5-RBD*, which was the least able to compete with wild-type Anillin for localization. That the double mutant showed even weaker localization than Anillin-RBD* suggests that the interaction with Sticky does provide a low-level contribution to equatorial Anillin localization. We conclude that the normal pattern of localization of Anillin at cleavage furrows reflects multiple pools of Anillin molecules engaged in distinct interactions including, but not limited to, Sticky and Rho1. Anillin is likely ordinarily dynamically switching between these pools to coordinate CR closure and MR formation.

## Discussion

Cytokinesis is a complex and dynamic process involving multiple cytoskeletal networks that cooperate in poorly understood ways to remodel the plasma membrane and generate two cells from one. The CR has been the accepted organelle of cell cleavage for over 50 years (Schroeder, 1972). However, there remains a significant lack of understanding as to how CRs assemble, generate tension, constrict while disassembling, and ultimately mature into stable MRs (Pollard, 2017). Anillin orthologues are of central importance and able to bind a host of other essential cytokinesis regulators, including F-actin, Myosin II, RhoA, septins, Citron kinase, importins, RacGAP50C, among others (Piekny and Maddox, 2010). Anillin is generally considered a “scaffold” that crosslinks components to the actomyosin ring. However, neither the functional roles of its diverse interactions nor their coordination with one another are well understood. Optically resolving different cytoskeletal elements within the CR and MR remains a major challenge because the components become highly compacted in a diffraction-limited manner and undergo rapid turnover. However, by exploiting a simple assay in which different separation-of-function alleles of Anillin are co-expressed in the same *Drosophila* S2 cells, we find that separate pools of Anillin, scaffolding distinct cytoskeletal elements, promote successful cytokinesis. This provides compelling evidence, as proposed in (Carim et al., 2020), that cytokinetic ring structures contain cytoskeletal networks that are not just coupled but also *uncoupled* from one another and thus free to move apart, while being independently anchored to the plasma membrane. This important finding can help explain the long-standing enigmas of how CRs close and how MRs form.

We first define a novel separation-of-function allele, Anillin-Δ1-5, that lacks the first 5 conserved amino acids and that loses the ability to interact with the Citron kinase, Sticky. Anillin-Δ1-5 failed to support a faithful CR-to-MR transition in deplete-and-rescue experiments but was nonetheless able to complement Anillin-RBD*, another allele previously shown to be defective in Rho1-GTP binding, anillo-septin assembly and CR closure (Carim and Hickson, 2023). The functional trans-heterozygous combination of Anillin-Δ1-5 and Anillin-RBD* demonstrates that the opposing actions of Sticky and septins on Anillin, required to mediate the transition from CR to MR (El Amine et al., 2013), do not need to occur simultaneously on the same Anillin molecules. Rather, different Anillin molecules capable of interacting with either Sticky or with Rho1/septins can functionally complement one another. It has been suggested that the unstructured N-terminus of *C. elegans* ANI-1 might oligomerize (Lebedev et al., 2023). However, unless oligomerization requires the first five amino acids and is not required for function, our observations of *Drosophila* Anillin do not support this prediction, since the two co-expressed Anillin species localized independently under all conditions, including in LatA.

We previously concluded that Anillin acts as a “bifunctional linker” based on the behaviors of its two halves (Kechad et al., 2012). However, the present study suggests that Anillin can engage with subsets of interactors in a mutually exclusive manner and that these are not linked. This calls for a revised model for the CR-to-MR transition (Fig. 6), in which the balance between retention versus removal of ring material involves the physical separation of distinct, membrane-anchored cytoskeletal complexes, each scaffolded by different molecules of Anillin. Anillin also binds and bundles F-actin (Field and Alberts, 1995; Jananji et al., 2017; Tian et al., 2015), which can generate contractile force even in the absence of myosin (Kučera et al., 2021). F-actin is lost during the transition from CR to MR, and this transition is severely disrupted upon deletion of the actin binding domain of Anillin (Jananji et al., 2017). We therefore propose that depolymerization of Anillin-scaffolded F-actin during CR closure locally releases Anillin molecules, which then engage either with Rho1-GTP (and subsequently the septins), or with Citron kinase/Sticky. Anillin can also bind the myosin II regulatory light chain (MRLC) via a conserved myosin binding domain which sits between the F-actin binding domain and the Sticky-interacting NTD (Straight et al., 2005). Although the functional consequence of the MRLC-Anillin interaction remains unclear, we note that MRLC (Spaghetti squash in *Drosophila*) remains a prominent marker of the mature MR, along with Sticky and Anillin (El Amine et al., 2013; El-Amine et al., 2019; Kechad et al., 2012). It therefore seems reasonable to propose that Anillin molecules may simultaneously bind MRLC and Sticky during CR disassembly and MR formation. Retention of Anillin at the MR likely also requires interactions between Sticky or Anillin and microtubule associated proteins at the interdigitating microtubules of the midbody (Bassi et al., 2013; D’Avino et al., 2008; McKenzie et al., 2016; Watanabe et al., 2013). Conversely, removal from the nascent MR involves Rho1-GTP-dependent anillo-septin assembly and shedding (Carim and Hickson, 2023; El Amine et al., 2013). This model satisfactorily accounts for the observed rescue and physical separation of Anillin-Δ1-5 and Anillin-RBD* in our trans-heterozygous combination experiments (see Fig. S4 for model).

**Fig. 6.**
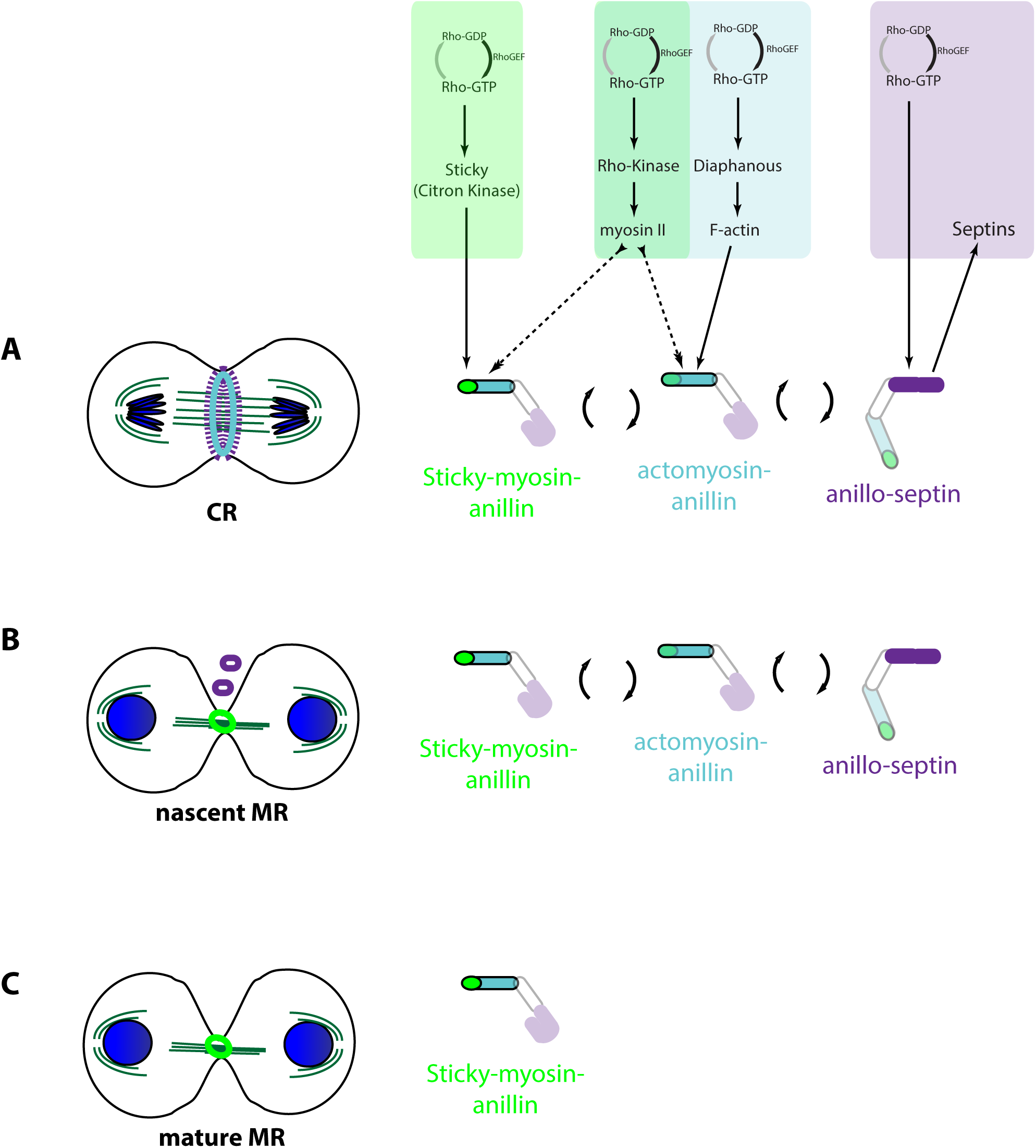
Model for how Anillin independently scaffolds distinct Rho1-dependent proteins to control the CR-to-MR transition. **A** At the CR, Anillin molecules independently and dynamically scaffold actomyosin, Sticky- myosin, or Rho1/septins (anillo-septin) to coordinate CR closure with its disassembly. **B** At the nascent MR, actin disassembles, while Anillin molecules form either Sticky-myosin-anillin complexes that are retained in the nascent MR (light green), or anillo-septin complexes that are removed/shed (purple). **C** Upon complete disassembly of F-actin and removal of Rho1- dependent anillo-septin complexes, Rho1-dependent Sticky-myosin-anillin complexes are retained and incorporated into the mature MR.

It is important to highlight that all these events are coordinated by Rho1-GTP, which is continually generated at the equatorial plasma membrane during cytokinesis (Bement et al., 2005), by the RhoGEF Pebble/ECT2 (Somers and Saint, 2003), and which undergoes GTPase flux (Miller et al., 2008). Both sets of Anillin-scaffolded complexes described in the present study must be controlled by discrete pools of Rho1, since Rho1-GTP-bound Anillin is specifically required for anillo-septin assembly (and thus shedding) (Carim and Hickson, 2023), and Rho1-GTP-bound Citron kinase/Sticky is specifically required for MR formation (Eda et al., 2001; El-Amine et al., 2019; Watanabe et al., 2013). We provide evidence that both of these Rho1-dependent pathways contribute to Anillin localization. While the Anillin RBD-dependent pathway appears to be the major mechanism of localization at the CR stage, this may reflect the fact that the C-terminus of Anillin participates in a positive feedforward loop: Rho1-GTP recruits Anillin (and septins (Carim and Hickson, 2023)), which in turn prolongs the lifetime of the Rho1-GTP, presumably by preventing its GAP-stimulated hydrolysis (Budnar et al., 2019). Thus, the Anillin C-terminus may promote the accumulation of a reservoir of Pebble/ECT2-generated active Rho1 at the furrow, as first shown in (Piekny and Glotzer, 2008), which in turn sustains a pool of Anillin and septins at the plasma membrane. Conversely, distinct Rho1-GTP-Sticky-Anillin complexes may turn over more rapidly at the CR, through Rho1 GTPase flux, resulting in fewer Rho1-GTP-Sticky-Anillin complexes being present at any given time. We speculate that this underlies the relatively ‘weak’ localization observed for Anillin-RBD* at the CR (see Fig. S4). We further speculate that the observed accumulation and retention of Anillin-RBD* complexes at the later nascent MR, involves another mechanism preventing GTP hydrolysis of the Sticky-bound Rho1 that occurs once the appropriate conditions for the transition to MR formation are met (e.g. bundling of midbody microtubules). However, we note that the observed localization of Anillin-Δ1-5- RBD* to the midbody indicates the existence of additional mechanisms, independent of both Sticky-binding and Rho1-GTP binding, that must also contribute to Anillin localization at this stage.

Rho1-GTP also binds the formin, Diaphanous, and Rho-kinase, which mediate F-actin and myosin II recruitment to the CR. These proteins have not been studied here but likely also contribute to Anillin localization. Such contributions may mediate the localization of the Anillin-Δ1-5-RBD* double mutant and must also be integral to the mechanisms controlling closure of the CR and its transition to the MR. The current work clearly establishes the co- existence of separate functional pools of Anillin and Rho1 at the cell cortex. However, further work is evidently needed to elucidate how all pools of Rho1-GTP simultaneously and/or sequentially engage its multiple effectors (Diaphanous, Rho-kinase, Sticky and Anillin), how GTPase flux is controlled for each of these pools, and how Anillin molecules subsequently engage with and/or disengage from components of each of those effector pathways to promote the seamless progression of cytokinesis, from assembly of the CR to the formation of the mature MR.

### Materials and Methods Cloning and molecular biology

Sticky-mCherry and Sticky-miniCC2a constructs were described in (El-Amine et al., 2019). mCherry-tubulin was expressed from pAc-mCh-Tub (DGRC Stock 1462 ; https://dgrc.bio.indiana.edu//stock/1462 ; RRID:DGRC_1462). Codon-optimized, RNAi-resistant Anillin-GFP and Anillin-RBD*-GFP (specifically Anillin-A874D-E892K-GFP) were described in (Carim and Hickson, 2023). Anillin-Δ1-5, and Anillin-Δ1-5-RBD* were PCR amplified from these, cloned into pENTR™/D-TOPO® (Invitrogen/ThermoFisher Scientific #k2400-20), sequence-verified and recombined into metallothionein promoter-driven pMT- WG (C-terminal GFP tag, Terence Murphy collection) and pMT-WCh (C-terminal mCherry tag) destination plasmids using LR clonase™ (Gateway® Invitrogen/Thermo Fisher Scientific #11791-019). DNA constructs were propagated in DH5α bacteria grown in Luria-Bertani (LB) broth supplemented with ampicillin and purified using QIAprep® Spin Miniprep kits (Qiagen #27106). DNA concentration was measured with a Nanodrop™ spectrophotometer, and plasmids were stored at -20°C prior to transfection.

### S2 Cell culture and transfections

Drosophila Schneider 2 (S2) cells (obtained from University of San Francisco, California) were cultured in Schneider’s Drosophila medium (Gibco/Thermo Fisher Scientific #21720- 024) supplemented with 10% fetal bovine serum (FBS, Invitrogen/Thermo Fisher Scientific #16140-071) and 3% penicillin-streptomycin (Invitrogen/Thermo Fisher Scientific #15140- 122), at 25°C in ambient CO_2_. Cells were routinely passaged every 7 days. 7.5 x 10^5^ cells/mL cells were plated in 12 well culture plates and transiently transfected 24 h after passaging. 1.0μg of each DNA plasmid was mixed in 100μL serum-free Schneider’s Drosophila medium with FugeneHD (Promega, #E2311), using a 3:1 Fugene:DNA ratio. Fugene-DNA mixes were incubated for 20 min at room temperature before being adding to cells in complete medium.

### RNAi and drug treatments

Double-stranded RNAs (dsRNA) were generated in a two-step PCR amplification using plasmid cDNA as the starting template. In the first step PCR, the following gene-specific primer pairs each containing an 8 bp anchor (5’-GGGCGGGT-3’) were used: LacI (5′- GGGCGGGTTGGTGGTGTCGATGGTAGAA-3′ and 5′-GGGCGGGTCGGTATCGTCGTATCCCACT- 3′); anillin dsRNA1 (5′-GGGCGGGTTAGAAATCTATGGCATGTTGGC-3′ and 5′- GGGCGGGTGAGAAAACTGTTAACAACCCGC-3′); sticky dsRNA1 (5′- GGGCGGGTGTGAAACCGTTGGTGATATGC-3′ and 5′-GGGCGGGTTTCAACCTCTGGAAGTTATCG-3′). The resulting PCR products were purified using QIAquick® PCR purification kit (Qiagen #28104) and used as templates for a second round of PCR using a universal T7 primer (5’-TAATACGACTCACTATAGGGAGACCACGGGCGGGT-3’) that anneals to the anchor sequence and incorporates the T7 promoter sequence. DsRNAs were generated by *in vitro* transcription overnight at 37°C using T7 RiboMAX™ kits (Promega #P1300). Template cDNA was then digested by RNase-free DNase RQ1 (Promega #M610C) for 15 min at 37°C. The remaining single strand RNAs were precipitated in ethanol, air dried, and resuspended in nuclease-free water, before being annealed in a 95°C water bath allowed to slowly cool to room temperature. Resulting dsRNAs were verified by agarose gel electrophoresis, quantified by spectrophotometry (Nanodrop) and stored at -20°C until use.

For dsRNA treatments, cells were seeded in complete medium in 96-well glass-bottomed imaging plates (SensoPlate™ VWR #30618-030) and incubated with 1 μg/mL of dsRNA for 36 to 72 h. For rescue experiments, CuSO4 (0.5 mM final concentration) was added at the same time as the dsRNAs to induce expression of the *metallothionein* promoter. For non-rescue experiments, CuSO_4_ induction was performed 16-24 h prior to imaging. Where applicable, cells were treated with Latrunculin A (LatA, EMD Millipore 701687) at a final concentration of 1μg/ml, 20-30 min before the start of image acquisition.

### Microscopy

Imaging was performed at 21°C using a spinning disk confocal microscope system: Leica DMI6000B stand with motorized stage with piezo-Z insert (ASI), Perkin Elmer® UltraVIEW VoX equipped with a charge-coupled device (CCD) camera (ORCA-R2 Hamamatsu). Image acquisition was controlled by Volocity version 6.3 (Perkin Elmer®) and performed in emission discrimination mode, using 488 nm and 561 nm lasers to excite GFP and mCherry, respectively. Camera binning was set to 2×2 and exposure times 200 ms, with laser power adjusted to ensure signals were within the dynamic range of the camera. Confocal stacks of 10-12 μm were collected, with an optical spacing of 1 µm. High-resolution imaging was performed for short-duration acquisitions (less than 3 hours) with images taken every 1-2 min using a plan apochromat 63× 1.4NA oil immersion objective. For scoring division attempt outcomes, lower resolution imaging was performed overnight using a plan apochromat 40×0.85 NA air objective, with images taken every 5 min.

### Image analysis, quantification and statistical analyses

Images acquired using Volocity version 6.3 (Quorum Technologies) were exported to Photoshop CS6 (Adobe) for assembling panels, and Figures were then produced in Illustrator CS6 (Adobe). Unless otherwise stated in the figure legends, displayed images are maximum intensity projections of multiple focal planes.

Ring diameter measurements were performed in Volocity using the line measurement tool at the central Z-plane and were normalized relative to the initial diameter at anaphase onset.

For measurements of cortical localization, datasets were exported to FIJI (Image J). A line was manually drawn along the cortex from pole to pole at each timepoint of interest and using the confocal section that best captured the center of the cell. For each line and for each channel, intensity values were corrected by subtracting background intensities determined from a line drawn outside the cell. For quantitation of the total cortical localization, the sum of all background-corrected intensity values along the line was calculated for each channel, and the sums from each time point was then normalized to the sums obtained at T=0 (anaphase onset). For measurements of equatorial cortex enrichment, each pole-to-pole line was first segmented into 4 equal lengths: the two outer segments were designated as “polar”; the two inner segments were designated as “equatorial”. Equatorial enrichment was then calculated for each channel as the ratio of the sums of “equatorial”: “polar” intensity values for that channel. For Fig 5E, where mutants were co-expressed with wild-type Anillin-GFP, intensities values were obtained from measurement lines across the cell equator, perpendicular to the spindle axis, and the mean peak cortex values were divided by the mean cytoplasm values (all background-corrected) to determine cortex: cytoplasm ratios. The cortex: cytoplasm ratios of the indicated mCherry-tagged constructs were then normalized to the corresponding cortex: cytoplasm ratio of wild-type Anillin-GFP co-expressed in the same cell.

Graphs and statistical analyses were performed in GraphPad Prism 7. Unpaired t-tests compared the means between two groups. Unpaired multiple t-tests involved comparing the mean at each indicated time between two conditions. The one-way ANOVA test (Kruskal-Wallis test, Fig. 4F) compared the mean of one group with the mean of each other group. P-values are displayed as follows: ns = p> 0.05; *, p<0.05, **, p < 0.01; *** p< 0.001; **** p < 0.0001. Unless otherwise specified, each experiment was repeated three times (n=3).

## Supplemental material

Fig. S1, related to Fig. 1, shows full blots of GST-AnillinΔ1-5-NTD pulldowns and co-expression of Anillin or AnillinΔ1-5 with Sticky. Fig. S2, related to Fig. 4, shows phenotypic characterization of the AnillinΔ1-5-RBD* cis double mutant. Fig. S3, related to Fig. 4, shows the characterization of trans-heterozygous expression of AnillinΔ1-5 and Anillin-RBD* with GFP and mCherry tags swapped. Fig. S4, related to Fig. 6, shows a cartoon model for how trans-heterozygous expression of AnillinΔ1-5 and Anillin-RBD* provides function. Video 1 is related to Fig. 1; Video 2 is related Fig. 2; Videos 3 and 4 are related to Fig. 3; Videos 5, 6 and 7 are related to Fig. 4; Videos 8 and 9 are related to Fig. 5.

## Data availability statement

Data are available in the primary article and in the supplementary materials. Original data and plasmids generated in this study are available upon request from the corresponding author.

## Acknowledgements

We thank members of the Hickson lab, especially Amel Kechad, for discussions. We acknowledge the Drosophila Genomics Resource Center (NIH Grant 2P40OD010949). This research was supported by grants from the Canadian Institutes for Health Research (CIHR, reference PJT-183759), the Natural Sciences and Engineering Research Council (NSERC, reference RGPIN-2021-03324) and the Canadian Fund for Innovation (CFI) awarded to G.R.X.H. G.C received studentship funding from the Programme de Biologie Moléculaire and the Faculté de Médecine, Université de Montréal. S.C.C. received a postdoctoral fellowship from the Foundation CHU Sainte-Justine. The authors declare no competing financial interests.

## Author contributions

S. Carim and G. Hickson conceptualized the project. S. Carim generated and validated the novel Anillin-Δ1-5 separation-of-function mutant. G. Chambaud designed and conducted the co-expression experiments and performed their formal analysis. G. Chambaud and G. Hickson designed the figures. G. Hickson wrote the manuscript, incorporating feedback from all authors. G. Hickson obtained funding and supervised the project.

## Abbreviations

CR: contractile ring
LatA: latrunculin A
MR: midbody ring
NTD: N-terminal domain.

## Video legends

Video 1. **Drosophila S2 cell co-expressing Anillin-Δ1-5-GFP (green) and mCherry-tubulin (magenta) following endogenous anillin depletion.** Related to Fig. 1. Image stacks were acquired every 120 s on an UltraView Vox spinning disc confocal microscope (PerkinElmer) attached to a DMI600B stand (Leica Microsystems), equipped with a 63x 1.4- NA Plan-Apochromat oil immersion objective and ORCA-R2 CCD camera (Hamamatsu). Maximum intensity projection of the image stack is shown. Video starts at anaphase onset. Playback rate is 600x real time (5 frames/s).

Video 2. **Drosophila S2 cell co-expressing wild-type Anillin-GFP (green) and Anillin- mCherry (magenta) following endogenous anillin depletion.** Related to Fig. 2. Images were acquired every 60 s on an UltraView Vox spinning disc confocal microscope (PerkinElmer) attached to a DMI600B stand (Leica Microsystems), equipped with a 63x 1.4- NA Plan-Apochromat oil immersion objective and ORCA-R2 CCD camera (Hamamatsu). Maximum intensity projection of the image stack is shown. Video starts at anaphase onset. Playback rate is 300x real time (5 frames/s).

Video 3. **Drosophila S2 cell co-expressing AnillinΔ1-5-GFP (green) and AnillinΔ1-5- mCherry (magenta) following endogenous anillin depletion.** Related to Fig. 3. Images were acquired every 120 s on an UltraView Vox spinning disc confocal microscope (PerkinElmer) attached to a DMI600B stand (Leica Microsystems), equipped with a 63x 1.4- NA Plan-Apochromat oil immersion objective and ORCA-R2 CCD camera (Hamamatsu). Maximum intensity projection of the image stack is shown. Video starts at anaphase onset. Playback rate is 600x real time (5 frames/s).

Video 4. **Drosophila S2 cell co-expressing Anillin-RBD*-GFP (green) and Anillin-RBD*- mCherry (magenta) following endogenous anillin depletion.** Related to Fig. 3. Images were acquired every 120 s on an UltraView Vox spinning disc confocal microscope (PerkinElmer) attached to a DMI600B stand (Leica Microsystems), equipped with a 63x 1.4- NA Plan-Apochromat oil immersion objective and ORCA-R2 CCD camera (Hamamatsu). Maximum intensity projection of the image stack is shown. Video starts at anaphase onset. Playback rate is 600x real time (5 frames/s).

Video 5. **Drosophila S2 cell co-expressing AnillinΔ1-5-RBD*-GFP (green) and AnillinΔ1- 5-RBD*-mCherry (magenta) following endogenous anillin depletion.** Related to Fig. 4. Images were acquired every 120 s on an UltraView Vox spinning disc confocal microscope (PerkinElmer) attached to a DMI600B stand (Leica Microsystems), equipped with a 63x 1.4- NA Plan-Apochromat oil immersion objective and ORCA-R2 CCD camera (Hamamatsu). Maximum intensity projection of the image stack is shown. Video starts at anaphase onset. Playback rate is 600x real time (5 frames/s).

Video 6. **Drosophila S2 cell co-expressing Anillin-Δ1-5-GFP (green) and Anillin-RBD*- mCherry (magenta) following endogenous anillin depletion.** Related to Fig. 4. Images were acquired every 120 s on an UltraView Vox spinning disc confocal microscope (PerkinElmer) attached to a DMI600B stand (Leica Microsystems), equipped with a 63x 1.4- NA Plan-Apochromat oil immersion objective and ORCA-R2 CCD camera (Hamamatsu). Maximum intensity projection of the image stack is shown. Video starts at anaphase onset. Playback rate is 600x real time (5 frames/s).

Video 7. *Drosophila* S2 cell co-expressing Anillin-Δ1-5-GFP (green) and Anillin-RBD*- mCherry (magenta), depleted of endogenous Anillin and pre-treated with 1 µg/ml LatA. Related to Fig. 4. Images were acquired every 120 s on an UltraView Vox spinning disc confocal microscope (PerkinElmer) attached to a DMI600B stand (Leica Microsystems), equipped with a 63x 1.4-NA Plan-Apochromat oil immersion objective and ORCA-R2 CCD camera (Hamamatsu). Maximum intensity projection of the image stack is shown. Video starts 15 min before anaphase onset. Playback rate is 600x real time (5 frames/s).

Video 8. ***Drosophila* S2 cell co-expressing Anillin-GFP (green) and Anillin-Δ1-5- mCherry (magenta), depleted of endogenous Anillin.** Related to Fig. 5. Images were acquired every 120 s on an UltraView Vox spinning disc confocal microscope (PerkinElmer) attached to a DMI600B stand (Leica Microsystems), equipped with a 63x 1.4-NA Plan-Apochromat oil immersion objective and ORCA-R2 CCD camera (Hamamatsu). Maximum intensity projection of the image stack is shown. Video starts at anaphase onset. Playback rate is 600x real time (5 frames/s).

Video 9. ***Drosophila* S2 cell co-expressing Anillin-GFP (green) and AnillinRBD*-mCherry (magenta), depleted of endogenous Anillin.** Related to Fig. 5. Images were acquired every 120 s on an UltraView Vox spinning disc confocal microscope (PerkinElmer) attached to a DMI600B stand (Leica Microsystems), equipped with a 63x 1.4-NA Plan-Apochromat oil immersion objective and ORCA-R2 CCD camera (Hamamatsu). Maximum intensity projection of the image stack is shown. Video starts at anaphase onset. Playback rate is 600x real time (5 frames/s).

