## Supplemental Figures and Legends for "Heteroallelic combination of anillin mutants reveals cytoskeletal uncoupling during cytokinesis"

### Supplementary Figures and Legends

**Figure S1. Residues 1-5 of Anillin are required to interact with Sticky and promote Sticky localization at the nascent MR.** Related to Fig. 1. **A** Alphafold2 predicted structure of Anillin/Scraps, showing N-terminal alpha-helix (red oval) and the position of residues 1-5 that are deleted in Anillin-Δ1-5. **B-C** Full blots of GST-Anillin-NTD (B) and GST-AnillinΔ1-5-NTD (C) pulldowns of Sticky-miniCC2a-GFP from whole S2 cell extract (WCE), immunoblotted with anti-GFP antibody (upper panels), including Ponceau-S staining of proteins prior to immunoblotting (lower panels). P, pellet; S, supernatant. **D** Stills from a representative time-lapse sequence of S2 cell transiently co-expressing Anillin-mCherry and Sticky-GFP undergoing cytokinesis, following depletion of endogenous Anillin. **E** Quantification of equatorial cortex enrichment of co-expressed Anillin-mCherry and Sticky-GFP, calculated as the fold increase of sum intensity at the equatorial cortex (defined as the two central 25% lengths of each measurement line) relative to the sum intensity at the polar cortex (defined as the two outer 25% lengths of each measurement line), mean ± SD from N=17 cells. **F** Stills from a representative time-lapse sequence of S2 cell transiently co-expressing AnillinΔ1-5-mCherry and Sticky-GFP undergoing cytokinesis, following depletion of endogenous Anillin. **G** Quantification of equatorial cortex enrichment of co-expressed Anillin-Δ1-5-mCherry and Sticky-GFP, calculated as the fold increase of sum intensity at the equatorial cortex (defined as the two central 25% lengths of each measurement line) relative to the sum intensity at the polar cortex (defined as the two outer 25% lengths of each measurement line), mean ± SD from N=15 cells. Times are h:min:s from anaphase onset; scale bars, 5 µm.

Fig. S1

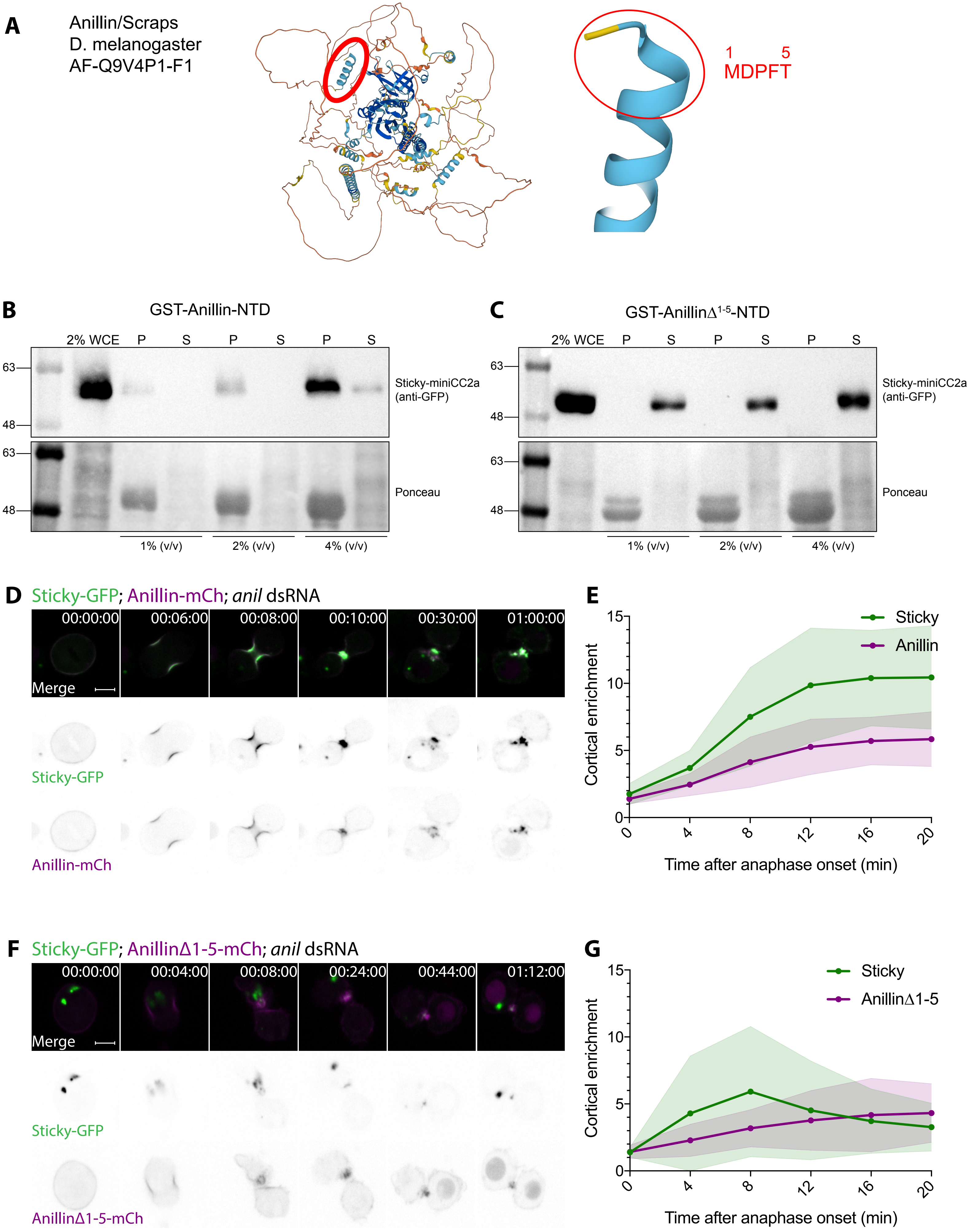
A
Anillin/Scraps
D. melanogaster
AF-Q9V4P1-F1
1 5
MDPFT
B
GST-Anillin-NTD
2% WCE P S P S P S
63
48
Sticky-miniCC2a
(anti-GFP)
Ponceau
1% (v/v)
2% (v/v)
4% (v/v)
C
GST-AnillinΔ1-5-NTD
2% WCE P S P S P S
Sticky-miniCC2a
(anti-GFP)
Ponceau
1% (v/v)
2% (v/v)
4% (v/v)
D
Sticky-GFP; Anillin-mCh; anil dsRNA
00:00:00 00:06:00 00:08:00 00:10:00 00:30:00 01:00:00
Merge
Sticky-GFP
Anillin-mCh
E
Sticky
Anillin
Cortical enrichment
Time after anaphase onset (min)
F
Sticky-GFP; AnillinΔ1-5-mCh; anil dsRNA
00:00:00 00:04:00 00:08:00 00:24:00 00:44:00 01:12:00
Merge
Sticky-GFP
AnillinΔ1-5-mCh
G
Sticky
AnillinΔ1-5
Cortical enrichment
Time after anaphase onset (min)

**Figure S2. Phenotypic characterization of the AnillinΔ1-5-RBD* cis double mutant.** Related to Fig. 4. **A** Outcomes of division attempts, scored through time lapse spinning disc confocal microscopy, of *Drosophila* S2 cells transiently co-expressing AnillinΔ1-5-RBD*-mCherry and AnillinΔ1-5-RBD*-GFP, following 3-day depletion of endogenous Anillin. Timing of stated outcome is relative to anaphase onset. Note that "failure" corresponds to observed furrow regression and binucleation, while "no failure" can signify either successful abscission or the end of the recording, and thus may include cells that were destined to fail. Mean ± SD from 122 division attempts are shown, ****: p-value < 0.0001. **B** Cortex: cytoplasm ratios of co-expressed AnillinΔ1-5-RBD*-mCherry and AnillinΔ1-5-RBD*-GFP measured from linescans across the equator during cytokinesis. Mean ± SD from N=12 cells, n=2 experiments are shown. **C** Cortex: cytoplasm ratios of co-expressed AnillinΔ1-5-RBD*-mCherry and wild-type Anillin-GFP measured from linescans across the equator during cytokinesis. Mean ± SD from N=22 cells are shown. **D** Quantification of equatorial cortex enrichment of co-expressed AnillinΔ1-5-RBD*-mCherry and wild-type Anillin-GFP, calculated as the fold increase of sum intensity at the equatorial cortex (defined as the two central 25% lengths of each measurement line) relative to the sum intensity at the polar cortex (defined as the two outer 25% lengths of each measurement line), mean ± SD from N=22 cells are shown.

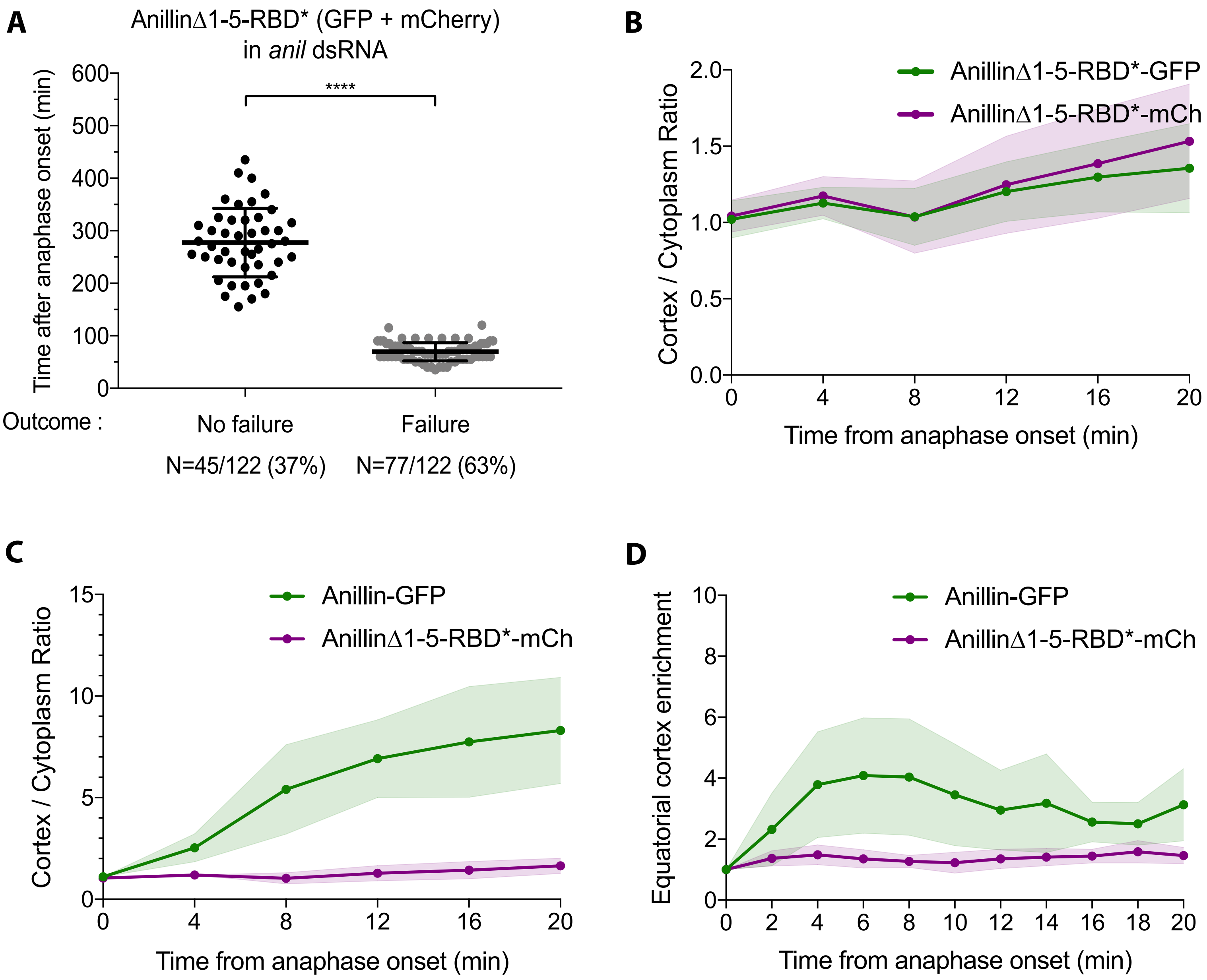
A
AnillinΔ1-5-RBD* (GFP + mCherry)
in *anil* dsRNA
Time after anaphase onset (min)
0
100
200
300
400
500
600
****
Outcome :
No failure
Failure
N=45/122 (37%)
N=77/122 (63%)
B
Cortex / Cytoplasm Ratio
0.0
0.5
1.0
1.5
2.0
AnillinΔ1-5-RBD*-GFP
AnillinΔ1-5-RBD*-mCh
0
4
8
12
16
20
Time from anaphase onset (min)
C
Cortex / Cytoplasm Ratio
0
5
10
15
Anillin-GFP
AnillinΔ1-5-RBD*-mCh
0
4
8
12
16
20
Time from anaphase onset (min)
D
Equatorial cortex enrichment
0
2
4
6
8
10
Anillin-GFP
AnillinΔ1-5-RBD*-mCh
0
2
4
6
8
10
12
14
16
18
20
Time from anaphase onset (min)

**Figure S3. Characterization of trans-heterozygous expression of AnillinΔ1-5 and Anillin-RBD* with GFP and mCherry tags swapped.** Related to Fig. 4. **A** Stills from a representative time-lapse sequence of Drosophila S2 cell transiently co-expressing Anillin-RBD*-GFP and AnillinΔ1-5-mCherry undergoing cytokinesis, following 3-day depletion of endogenous Anillin. **B** Quantification of equatorial cortex enrichment of co-expressed AnillinΔ1-5-mCherry and Anillin-RBD*-GFP, calculated as the fold increase of sum intensity at the equatorial cortex (defined as the two central 25% lengths of each measurement line) relative to the sum intensity at the polar cortex (defined as the two outer 25% lengths of each measurement line), mean ± SD from N=16 cells, n=2 experiments are shown. **C** Cortex: cytoplasm ratios of co-expressed AnillinΔ1-5-mCherry and Anillin-RBD*-GFP measured from linescans across the equator during cytokinesis. Mean ± SD from N=16 cells, n=2 experiments are shown **D** Quantification of equatorial cortex enrichment of co-expressed Anillin-RBD*-mCherry and AnillinΔ1-5-GFP, calculated as the fold increase of sum intensity at the equatorial cortex (defined as the two central 25% lengths of each measurement line) relative to the sum intensity at the polar cortex (defined as the two outer 25% lengths of each measurement line), mean ± SD from N=30 cells are shown. **E** Cortex: cytoplasm ratios of co-expressed Anillin-RBD*-mCherry and AnillinΔ1-5-GFP measured from linescans across the equator during cytokinesis. Mean ± SD from N=30 cells are shown.

Fig. S3

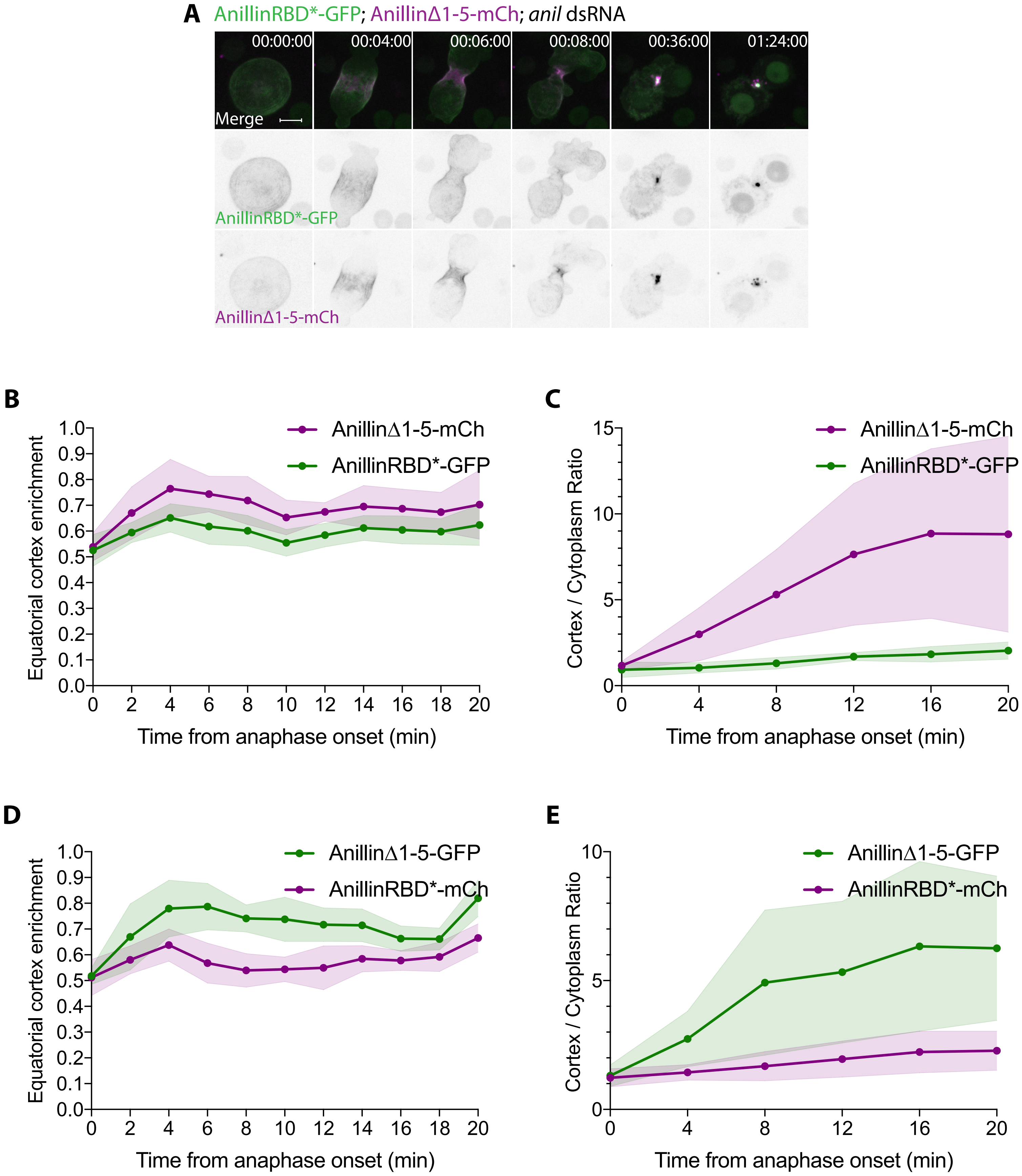

A
AnillinRBD*-GFP; AnillinΔ1-5-mCh; anil dsRNA
00:00:00
00:04:00
00:06:00
00:08:00
00:36:00
01:24:00
Merge
AnillinRBD*-GFP
AnillinΔ1-5-mCh
B
Equatorial cortex enrichment
Time from anaphase onset (min)
AnillinΔ1-5-mCh
AnillinRBD*-GFP
C
Cortex / Cytoplasm Ratio
Time from anaphase onset (min)
AnillinΔ1-5-mCh
AnillinRBD*-GFP
D
Equatorial cortex enrichment
Time from anaphase onset (min)
AnillinΔ1-5-GFP
AnillinRBD*-mCh
E
Cortex / Cytoplasm Ratio
Time from anaphase onset (min)
AnillinΔ1-5-GFP
AnillinRBD*-mCh

**Figure S4. Speculative model for how trans-heterozygous expression of AnillinΔ1-5 and Anillin-RBD* provides function.** Related to Fig. 6. Cartoon representations showing the abilities of both AnillinΔ1-5 and Anillin-RBD* to scaffold actomyosin, but each having the unique ability to associate with either Rho1-GTP-bound Sticky (Anillin-RBD*) or with Rho1-GTP/Septins (AnillinΔ1-5). We speculate that, at the CR stage, Sticky-myosin-anillin and actomyosin-anillin complexes undergo fast turnover, through Rho1 GTPase flux, while at the nascent MR stage, slowed turnover of Rho1-Sticky-myosin-anillin complexes leads to their retention. Conversely, we speculate that slow turnover of Rho1-GTP-dependent anillo-septin complexes occurs throughout, leading to the relative accumulation of these complexes at the CR and allowing their septin-dependent removal upon exclusion from the maturing MR.

Figure S4

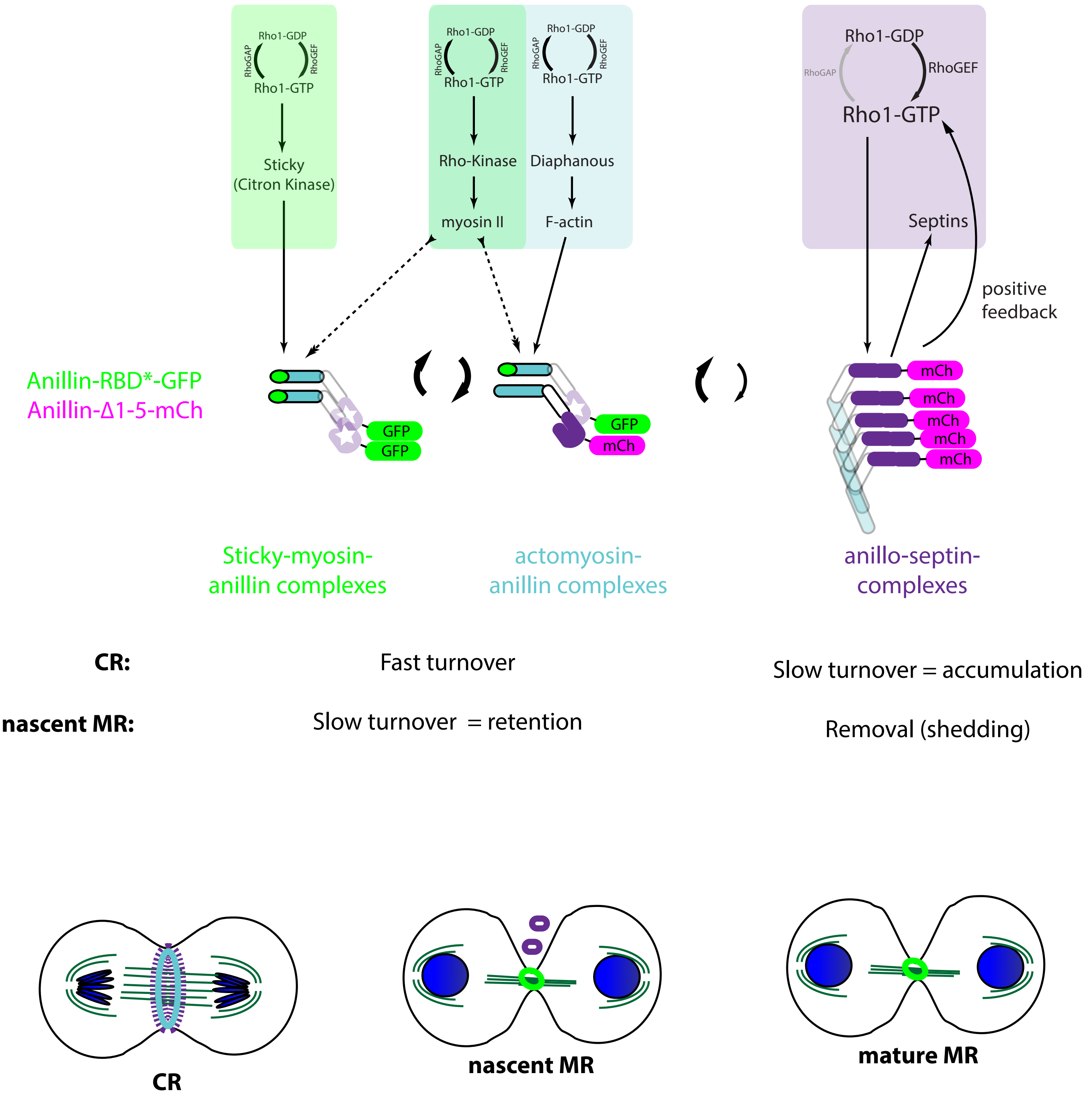
Rho1-GDP
RhoGAP
RhoGEF
Rho1-GTP
Sticky
(Citron Kinase)
Rho1-GDP
RhoGAP
RhoGEF
Rho1-GTP
Rho-Kinase
myosin II
Rho1-GDP
RhoGAP
RhoGEF
Rho1-GTP
Diaphanous
F-actin
Rho1-GDP
RhoGAP
RhoGEF
Rho1-GTP
Septins
positive
feedback
Anillin-RBD*-GFP
Anillin-Δ1-5-mCh
GFP
GFP
GFP
mCh
mCh
mCh
mCh
mCh
mCh
Sticky-myosin-
anillin complexes
actomyosin-
anillin complexes
anillo-septin-
complexes
CR:
Fast turnover
Slow turnover = accumulation
nascent MR:
Slow turnover = retention
Removal (shedding)
CR
nascent MR
mature MR
